# Erythromycin, Retapamulin, Pyridoxine, Folic acid and Ivermectin dose dependently inhibit cytopathic effect, Papain-like Protease and M^PRO^ of SARS-CoV-2

**DOI:** 10.1101/2022.12.28.522082

**Authors:** Shaibu Oricha Bello, Mustapha Umar Imam, Muhammad Bashir Bello, Abdulmajeed Yunusa, Adamu Ahmed Adamu, Abdulmalik Shuaibu, Ehimario Uche Igumbor, Zaiyad Garba Habib, Mustapha Ayodele Popoola, Chinwe Lucia Ochu, Aishatu Yahaya Bello, Yusuf Yahaya Deeni, Ifeoma Okoye

**Author notes:** Correspondence: Shaibu Oricha Bello, Department of Pharmacology and Therapeutics, Faculty of Basic Clinical Sciences, College of Health Sciences, Usmanu Danfodiyo University, Sokoto, Nigeria.

## Abstract

We previously showed that Erythromycin, Retapamulin, Pyridoxine, Folic acid and Ivermectin inhibit SARS-COV-2 induced cytopathic effect (CPE) in Vero cells. In this study and using validated quantitative neutral red assay, we show that the inhibition of CPE is concentration dependent with Inhibitory Concentration-50(IC_50_) of 3.27 μM, 4.23 μM, 9.29 μM, 3.19 μM and 84.31 μM respectively. Furthermore, Erythromycin, Retapamulin, Pyridoxine, Folic acid and Ivermectin dose dependently inhibit SARS-CoV-2 Papain-like Protease with IC_50_ of 0.94 μM, 0.88 μM, 1.14 μM, 1.07 μM, 1.51 μM respectively and the main protease(M^PRO^) with IC_50_ of 1.35 μM, 1.25 μM, 7.36 μM, 1.15 μM and 2.44 μM respectively. The IC_50_ for all the drugs, except ivermectin, are at the clinically achievable plasma concentration in human, which supports a possible role for the drugs in the management of COVID-19. The lack of inhibition of CPE by Ivermectin at clinical concentrations could be part of the explanation for its lack of effectiveness in clinical trials.

## 1. Introduction

A major lesson learned from the Spanish flu of 1918 is that pandemics can rapidly decimate a population^1^. In the current COVID-19 pandemic, which started in Wuhan China in December 2019, as of 13/11/2022, over 635million cases have been reported worldwide with over 6.61 Million deaths^2^. While the world continues to take measures aimed at keeping the incidence low and limiting the spread, it is important to identify therapeutics that may mitigate COVID-19 in those who do get infected and develop the disease. Although the world has risen to the challenge and there are over 400 drug candidates, most are based on new therapies that, if found effective, are not expected to be available in developing economies within 2 years due to market prioritization usually mostly imposed by limited production capacities of new drugs. Meanwhile, few candidate therapies have consistently been effective in ameliorating the duration of infection or the severity of the illness. Other than Dexamethasone and related steroid, Lopinavir-Ritonavir (Paxlovid) and Remdesivir are not available in most countries and are, nevertheless, effectively out of reach to most people due to cost. A huge therapeutic gap in COVID-19, therefore, remains.

COVID-19 is predominantly asymptomatic (80% of cases) ^3^. When symptomatic, the clinical presentation is that of respiratory infection with severity ranging from a mild common cold-like illness to severe viral pneumonia leading to acute respiratory distress syndrome that is potentially fatal ^3^. Knowledge of the pathogenesis of COVID-19 is still evolving but the structure of the virus, mechanisms of viral entry, and replications are essentially settled and appears target rich for drug discovery and development. It is, therefore, difficult to explain why many candidate therapies perform excellently well in-vitro but fail in clinical trials. The reason may include the presence of unresolved redundant pathways or a dis-linkage between viremia and pathology. Whichever, it is becoming clear that therapies that target multiple pathways in the pathogenesis of COVID-19 may prove superior.

Our team previously developed and validated an algorithm that involves deliberate consideration of multiple targets, cost, toxicity, and availability in selecting drugs for rapid repurposing efforts with off-label application as the immediate goal^4^. SARS-CoV-2 induced cytopathic effect(CPE) has previously been validated as a surrogate for SARS-CoV-2 infectivity and de-isolation which is non-inferior to PCR^5^. Therefore, in our previous study^4^ demonstration of inhibition of CPE was considered more cost-effective and a surrogate of clinical efficacy than demonstration of inhibition of viral multiplication alone due to the possibility of threshold effect in viral induced CPE^6^. We also identified Erythromycin, Pyridoxine, Folic acid and Ivermectin as potential drugs to repurpose for COVID-19; with wet laboratory demonstration of inhibition(qualitative assay) of SARS-COV-2 induced CPE in Vero cells and in-silico prediction of inhibition of multiple critical SARS-CoV-2 enzymes^4^.

This study was conducted to further evaluate Erythromycin, Pyridoxine, Folic acid and Ivermectin as potential drugs for COVID-19 by measuring quantitative concentration dependent inhibition of CPE, SARS-CoV-2 Papain-like Protease and SARS-CoV-2 3CL Main Protease(M^PRO^).

## 2. Materials and Methods

### 2.1 Materials

SensoLyte^®^ 520 SARS-CoV-2 M^PRO^ Activity Assay Kit, SensoLyte^®^ 520 SARS-CoV-2 Papain-like Protease/ Deubiq, Molecular grade water, Neutral red (3-amino-7-dimethylamino-2-methyl-phenazine hydrochloride) (Solarbio, cat. N8160), Minimal Essential Medium/Earls Balance Salts (MEM/EBSS) (HyClone laboratories, Utah, USA), Glacial acetic acid, Ethanol 96% 0.4% (wt./vol), Trypan blue in 0.9% NaCl solution, SARS-CoV-2, Virus (clinical isolate), and cell cultureware from Nest (Jiangsu, China) were used in this study.

### 2.2 Cell culture procedure

Vero cell preparation, passaging, SARS-CoV-2 virus, and sources of experimental drugs were as previously described^4^. Three different doses of each drug were tested for antiviral activity: Erythromycin, Retapamulin, Folic acid and Ivermectin were tested at 5μM, 7.5μM and 10μM, while Pyridoxine was tested at 10 μM, 15 μM and 20 μM, all in three independent replicates. Briefly, we seeded 96-well plates with 6×10^4^ cells/mL of Vero E6 (200 μL per well), using Minimum Essential Medium (MEM) with 10% fetal bovine serum (FBS) without antibiotics. Plates were incubated overnight at 37 °C in a 5% CO2 atmosphere. The following day, the 96 wells plates were viewed under an inverted microscope for a confluence of about 50%.

Sixty minutes before drug treatment, cell culture supernatant was removed from each well and the wells were washed with 150 μL phosphate buffered saline (PBS). Except for the negative and cell growth control wells where 50 μL of PBS was added, each well was infected with 50 μL SARS-CoV-2 diluted in PBS at a multiplicity of infection (MOI) of 0.1 and incubated for 1 hour at 37 °C in 5% CO2 with intermittent shaking of the plates at 15 minutes interval to allow for viral adsorption. Thereafter, the infection supernatant was removed and 200 μL of the respective drugs diluted in MEM/EBSS having 1% FBS without antibiotics were added to the different treatment groups and incubated at 37 °C in 5% CO2. The cells were viewed using an inverted microscope after 48 hours to check for CPE.

### 2.3 Quantification of Inhibition of SARS-COV-2 induced CPE using Neutral Red (NR) assay

Neutral red (NR) assay^7^ was used to quantify cell viability and the inhibition of CPE. The NR uptake assay provides a quantitative estimation of the number of viable cells in culture. It is based on the ability of viable cells to incorporate and bind the neutral red dye in the lysosomes which are then extracted using glacial acid for measurement of optical densities using a spectrophotometer^8^.

Briefly, an overnight incubated 40 μg ml^-1^ NR working solution (in MEM) was filtered using a 2 μm membrane filter to remove any precipitated dye crystals. The attached cells from the in vitro antiviral activity experiments were washed with 150 μL PBS per well and the washing solution was removed. The NR medium was gently placed in a reservoir and 100 μL of the NR medium was pipetted to each well of the plates. The plates were incubated for 2 h under the proper culture conditions. After the period of incubation, the plates were inspected with an inverted microscope to check the possible precipitation of NR. The medium was removed, and the cells were washed 2 times with 150 μL PBS per well. Thereafter 150 μL NR destain solution was added per well and the plates were shaken rapidly on a microtiter plate shaker at 500 rpm for 10 min.

The optical densities of extracted neutral red were measured at 540 nm in a microtiter plate reader. Blank was subtracted from the resulting absorbance values prior to data analysis. The groups were (i) Virus infected cells(virus control), (ii) Virus infected and treated cells(Treatment group) and (iii) Cells with no virus nor drug treatment(Growth control). Growth and inhibition of CPE was determined relative to the growth the untreated control.

### 2.4 Inhibition of SARS-CoV-2 Papain-like protease activity

All working solutions were prepared according to manufacturer instructions.

The enzymatic reaction was set up by adding 10 μL per well of three respective concentrations of Erythromycin, Retapamulin, Folic acid, Ivermectin (2.5μM, 5μM and 10μM) and pyridoxine (5 μM, 10 μM and 20 μM). Dose spacing was designed to be within the achievable plasma concentration of the drugs. Each test was done in triplicate.

Briefly, 40 μL diluted enzyme working solution was added to each microplate wells. Simultaneously the following control wells were set up.

i. Positive control containing 40 μL of Papain-like protease without the test drugs.
ii. Inhibitor control containing 40 μL Papain-like protease and 10 μL of GRL0617.
iii. Two vehicle control containing 40 μL Papain-like protease and 10 μL vehicle used in delivering test compound (0.1% DMSO).
iv. Test compound control containing assay buffer and 10 μL test compound.
v. Substrate control containing assay buffer.
vi. Assay buffer was used to bring the total volume of all controls to 50 μL.

To each well, 50 μL of Papain-like protease, substrate solution was added, and the reagents were mixed by shaking the plate gently for 30 sec. the plates were then incubated away from direct light at 37°C for up to 60 min. The fluorescence intensity was measured at excitation and emission wavelength of 490 nm and 520 nm.

### 2.5 Inhibition of SARS-CoV-2, M^PRO^ activity

The experimental set up and the relevant controls were as with the Inhibition of SARS-CoV-2 Papain-like protease activity assay, except that the SARS-CoV-2 M^PRO^ enzyme and activity assay kit was used according to manufacturer’s protocol.

## 3.0. Data Analysis

Statistical software GraphPad Prism 9.4.1, NCSS 2022 and Microsoft excel were used for data analysis and Graphical representations. We checked data for outliers using the Tukey’s box-plot method (interquartile principle)^9^. However, outliers were included in the analysis of CPE inhibition because biological reasons may be involved^9^. Furthermore, we substituted outliers by the group average in the enzyme assays because precision is more likely involved^10^. Nonetheless, sensitivity analysis was conducted by including outliers in the enzyme assays. Percentage Inhibition of CPE was calculated from the optical densities according to Severson *et all*^11^: 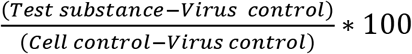, provided the concentrations of test drug was nontoxic to Vero cells in a separate assay. Although Severson *et al* considered inhibition of CPE ≥ 50% as hits in their study, we accepted inhibitions of CPE ≥ 15% as hits because this was not a full range dose-response study, but concentrations were restricted to those achievable within the human plasma at routine dosing schedules. Maximum responses (efficacy) were, therefore, conceivably different from what we observed. Percentage inhibition of enzyme activity was calculated according to manufacturer’s instruction and essentially represented as:

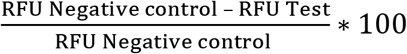 (after subtracting the blank), where RFU is the Relative Fluorescent Unit. The Concentration that gave 50% response(EC_50_ or IC_50_) was determined by first visual inspection of the dose response plot to infer possible models^12^. Maximum and minimum responses were constrained to 100% and 0% respectively because these were the only possible outcomes in data normalized to percentage response. Nonlinear regression three parameters dose-response models were used on the primary runs but compared to alternate models with higher R-square used to decide the best model. Furthermore, 95% Confidence Interval (CI) was determined for all estimates. In determining the 95% CI, unknowns were interpolated from standard curve. In the CPE evaluation all comparisons were to SARS-CoV-2 infected but vehicle (DMSO) treated Vero Cells. In the enzyme inhibition assay, all comparisons were to the uninhibited enzyme activity. In both enzyme inhibition assays, manufacturer supplied inhibitor controls were used to ascertain the integrity of the system. Level of significance was set at *p*<0.05.

## 4.0 Result

### 4.1 Inhibition of SARS-CoV-2 induced CPE

All tested doses of the drugs alone were not toxic to Vero cells.

Erythromycin, retapamulin, pyridoxine and folic acid dose dependently and significantly (p=0.0005) inhibited SARS-CoV-2 induced CPE with maximum inhibition of 76%, 75%, 63% and 42% respectively at the tested doses. However, the maximum inhibition of CPE by ivermectin was 10% (at 10 μM) and this was not significant (p=0.5)(Table 1 and Figure 1).

**Table 1:** Inhibition of SARS-CoV-2 induced CPE in Vero Cells

| Drug | Maximal inhibition of CPE(%) | Dose at maximal observed* inhibition of CPE($\mu\text{M}$) | IC<sub>50</sub>($\mu\text{M}$) | 95% CI of IC<sub>50</sub> ($\mu\text{M}$) | P |
| --- | --- | --- | --- | --- | --- |
| Erythromycin | 76 | 10 | 3.27 | 2.68 to 3.93 | <0.05 |
| Retapamulin | 75 | 7.5 | 4.23 | 2.29 to 7.22 | <0.05 |
| Pyridoxine | 63 | 20 | 9.29 | 6.30 to 13.31 | <0.05 |
| Folic Acid | 42 | 5 | 3.19 | 2.16 to 4.49 | <0.05 |
| Ivermectin | 10 | 7.5 | 84.31 | 41.70 to 759.2 | >0.05 |

\* 'Maximal observed' inhibition in this study is not the inherent maximum possible inhibition efficacy of the tested drug, but that observed within the dose range tested. These dose ranges were centred at achievable plasma concentration of the drugs at the current recommended doses except Ivermectin where ranges were derived from literature of concentrations previously found to inhibit SARS-CoV-2 proliferation<sup>13-15</sup>. Ivermectin cause non-significant inhibition of CPE and then at an IC<sub>50</sub> concentration not achievable in human plasma (800mcg/kg maximum single dose of ivermectin has $C_{\text{max}}$ 108.1ng/ml or 0.124 $\mu\text{M}/\text{ml}$ )<sup>16</sup>. Analyses were performed with outliers because biological difference probably dictated differences<sup>9</sup>.

**Figure 1:**
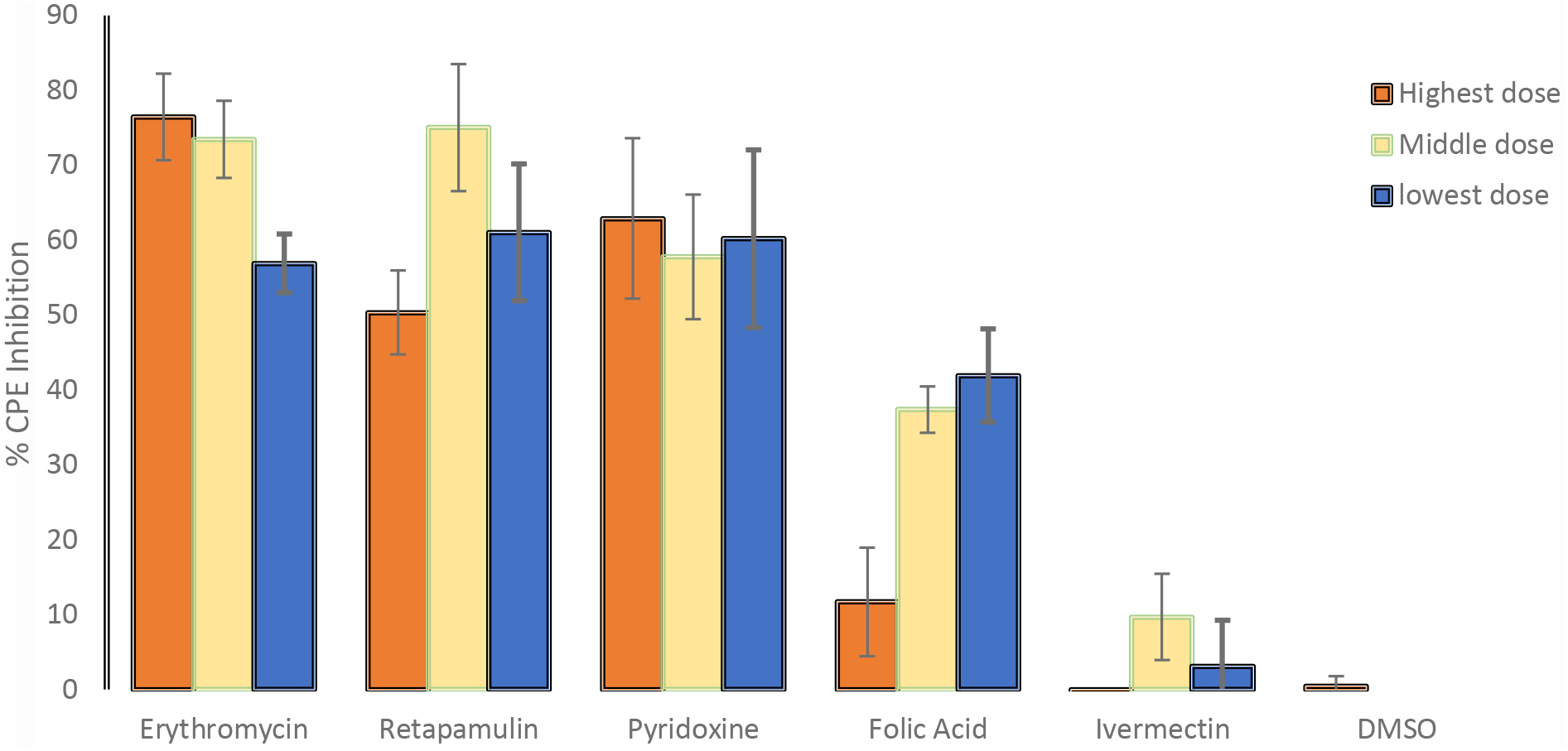
Inhibition of SARS-CoV-2 induced CPE. For Erythromycin, Retapamulin, Folic acid and Ivermectin high dose, medium dose and low dose respectively represent 10μM, 7.5μM and 5μM but for Pyridoxine the doses were 20 μM, 15 μM and 10 μM. The doses were selected based on achievable C_max_ in human subjects at standard doses of the drugs. All concentrations of test drugs were found nontoxic to Vero cells. Analyses were performed with outliers because biological factors probably dictated differences^9^.

### 4.2 Inhibition of SARS-CoV-2 Papain like Protease

Erythromycin, retapamulin, pyridoxine, folic acid and ivermectin dose dependently and significantly (p<0.05) inhibited SARS-CoV-2 papain like protease with maximum inhibition of 29.46%, 20.60%, 17.46%, 31.99% and 34.34% respectively at the tested doses (Table 2 and Figure 2).

**Table 2:** Inhibition of SARS-CoV-2 Papain like protease(PP)

| Drug | Maximal inhibition of PP(%) | Dose at maximal observed* inhibition of PP(μM) | IC<sub>50</sub>(μM) | 95% CI of IC<sub>50</sub> (μM) | p |
| --- | --- | --- | --- | --- | --- |
| Erythromycin | 29.46 | 10.0 | 0.94 | 0.20 to 2.09 | <0.05 |
| Retapamulin | 20.60 | 5 | 0.88 | 0.45 to 1.42 | <0.05 |
| Pyridoxine | 17.46 | 20.0 | 1.14 | 0.30 to 2.23 | <0.05 |
| Folic Acid | 31.99 | 5 | 1.07 | 0.47 to 1.90 | <0.05 |
| Ivermectin | 34.34 | 10 | 1.51 | 0.55 to 3.07 | <0.05 |

\* 'Maximal observed' inhibition in this study is not the inherent maximum possible inhibition efficacy of the tested drug, but that observed within the dose range tested. These dose ranges were centred at achievable plasma concentration of the drugs at the current recommended doses except Ivermectin where ranges were derived from literature of concentrations previously found to inhibit SARS-CoV-2 proliferation<sup>13-15</sup>. Ivermectin cause significant inhibition of PP but at an IC<sub>50</sub> concentration not achievable in human plasma (800mcg/kg maximum single dose of ivermectin has C<sub>max</sub> 108.1ng/ml or 0.124 μM/ml)<sup>16</sup>. Analyses were performed with substitution of outliers because precision rather than biological factors probably dictated differences<sup>9</sup>.

**Figure 2:**
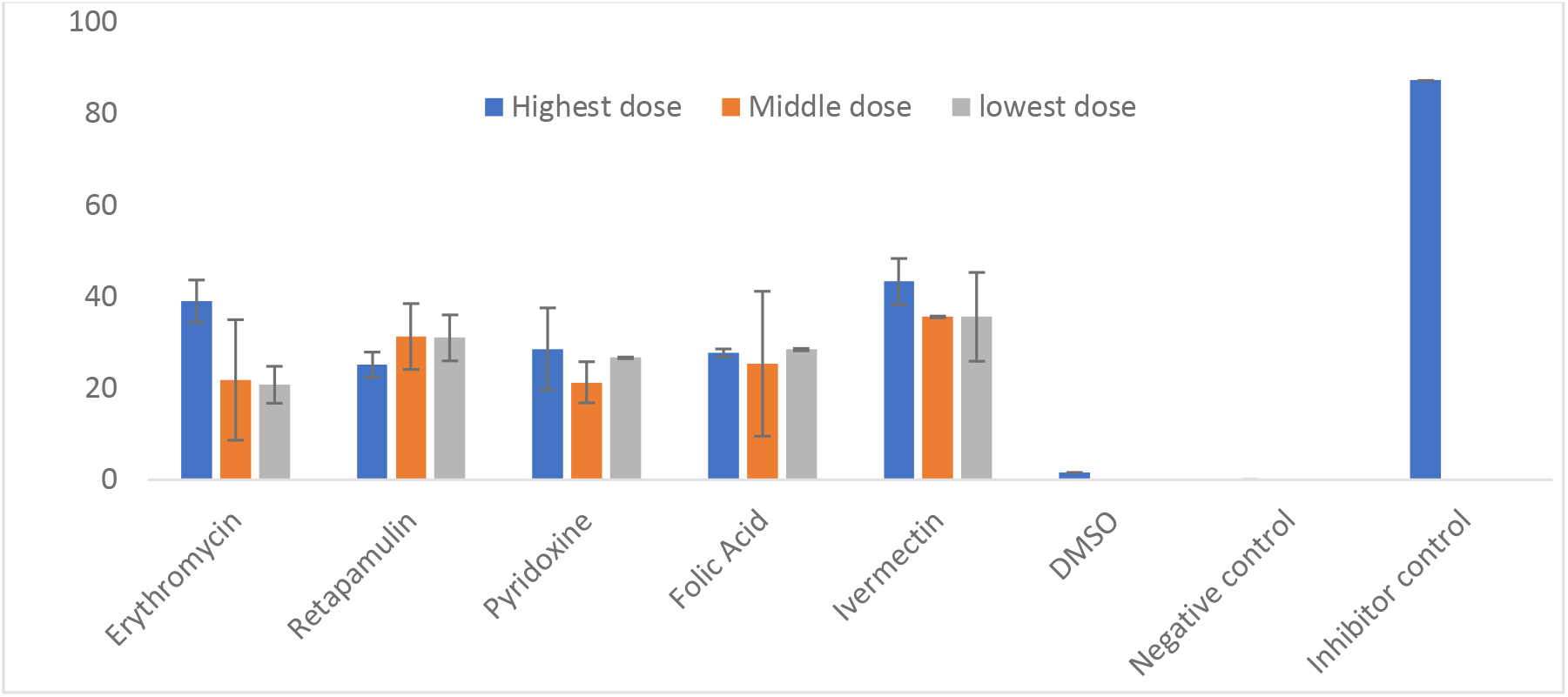
Inhibition of SARS-CoV-2 Papain like protease. For Erythromycin, Retapamulin, Folic acid and Ivermectin, high dose, medium dose and low dose respectively represent 10μM, 5μM and 2.5μM but for Pyridoxine the doses were 20 μM, 10 μM and 5 μM. The doses were selected based on achievable C_max_ in human subjects at standard doses of the drugs and to input dose doubling escalation method. All concentrations of test drugs were found nontoxic to Vero cells. Analyses were performed with outliers substituted by averaging or neighbourhood method because precision rather than biological factors probably dictated differences^9^.

### 4.3 Inhibition of SARS-CoV-2 Main Protease (M^PRO^ or 3CL protease)

Erythromycin, retapamulin, pyridoxine, folic acid and ivermectin dose dependently and significantly (p<0.05) inhibited SARS-CoV-2 M^PRO^ with maximum inhibition of 30.51%, 33.37%, 45.1%, 26.8% and 38.07% respectively at the tested doses (Table 3 and Figure 3).

**Table 3:** Inhibition of SARS-CoV-2 Main Protease(MPRO or 3CL)

| Table 3: Inhibition of SARS-CoV-2 Main Protease ($M^{PRO}$ or 3CL) | | | | | |
| --- | --- | --- | --- | --- | --- |
| Drug | Maximal inhibition of $M^{PRO}$ (%) | Dose at maximal observed* inhibition of $M^{PRO}$ ($\mu M$) | $IC_{50}$ ($\mu M$) | 95% CI of $IC_{50}$ ($\mu M$) | P |
| Erythromycin | 30.51 | 2.5 | 1.35 | 0.90 to 1.91 | <0.05 |
| Retapamulin | 33.37 | 2.5 | 1.25 | 0.60 to 2.15 | <0.05 |
| Pyridoxine | 45.17 | 20.0 | 7.36 | 4.14 to 12.67 | <0.05 |
| Folic Acid | 26.8 | 2.5 | 1.15 | 0.81 to 1.56 | <0.05 |
| Ivermectin | 38.07 | 5.0 | 2.44 | 1.49 to 3.77 | <0.05 |

\* 'Maximal observed' inhibition in this study is not the inherent maximum possible inhibition efficacy of the tested drug, but that observed within the dose range tested . These dose ranges were centred at achievable plasma concentration of the drugs at the current recommended doses except Ivermectin where ranges were derived from literature of concentrations previously found to inhibit SARS-CoV-2 proliferation<sup>13-15</sup> . Ivermectin cause significant inhibition of $M^{PRO}$ but at an $IC_{50}$ concentration not achievable in human plasma (800mcg/kg maximum single dose of ivermectin has $C_{max}$ 108.1ng/ml or 0.124 $\mu M$ /ml)<sup>16</sup>. Analyses were performed with substitution of outliers because precision rather than biological factors probably dictated differences<sup>9</sup>.

**Figure 3:**
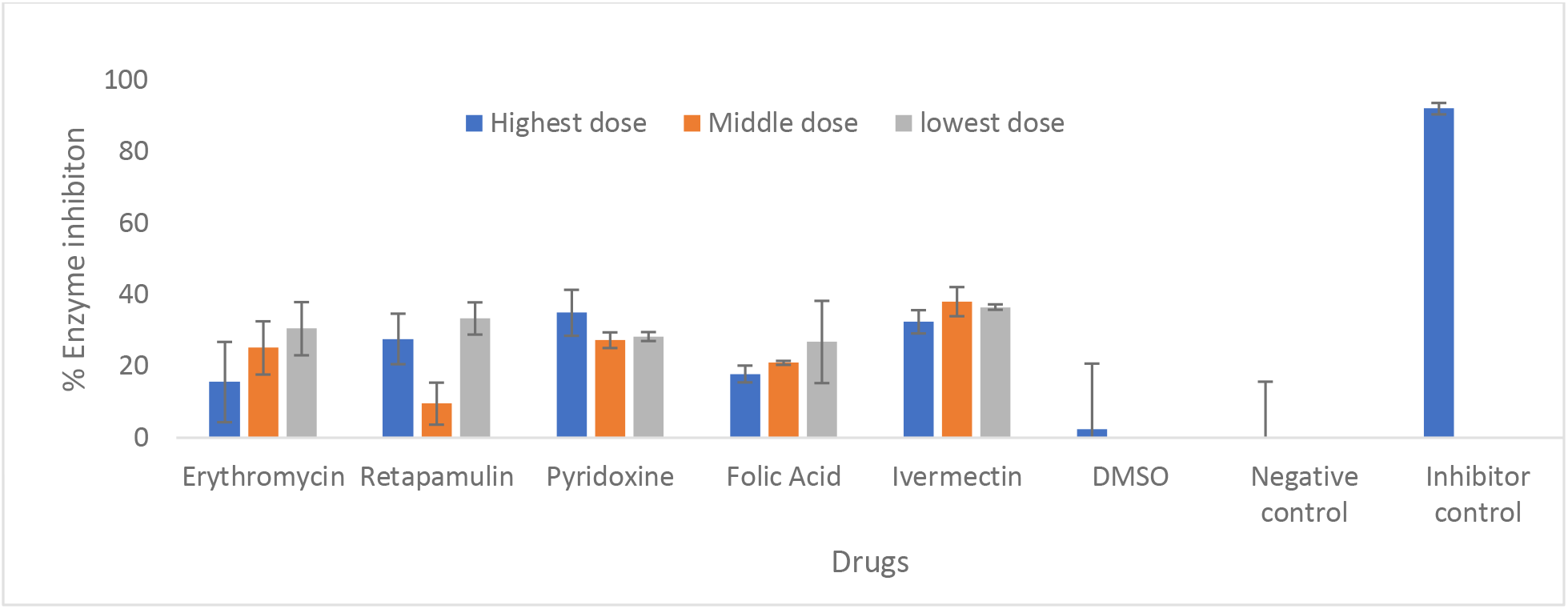
Inhibition of SARS-CoV-2 Main protease (MPRO or 3CL protease) For Erythromycin, Retapamulin, Folic acid and Ivermectin, high dose, medium dose and low dose respectively represent 10μM, 5μM and 2.5μM but for Pyridoxine the doses were 20 μM, 10 μM and 5 μM. The doses were selected based on achievable Cmax in human subjects at standard doses of the drugs and to input dose doubling escalation method. All concentrations of test drugs were found nontoxic to Vero cells. Analyses were performed with outliers substituted by averaging or neighbourhood method because precisions rather than biological factors probably dictated differences^9^.

## 5.0 Discussion

This study was conducted as standalone evaluation of the effect of drugs previously identified by in silico screening^4^ as potential therapeutics for SARS-CoV-2 infection and COVID-19. Although effect on SARS-CoV-2 induced CPE in Vero cells, SARS-CoV-2 Main protease (M^PRO^) and Papain-like protease (PP) were evaluated, these were not considered necessarily linked (mechanistic) because the initial in silico selection was based on predicted activity at 11 independent targets^4^, any combination of which may be related to observed inhibition of CPE. It is interesting that Nirmatrelvir-ritonavir combination(Paxlovid)^17,18^, the current forerunner in COVID-19 therapeutics is an M^PRO^ inhibitor and it is accepted that its efficacy is related to this inhibition^19^.

Erythromycin, retapamulin and pyridoxine significantly inhibited SARS-CoV-2 induced CPE in Vero cells to an extent that met our criteria (based on anticipated combination therapy) (≥15%) and that of Severson *et al* (based on anticipated monotherapy) (≥50%)^11^ to define hits. It is reassuring that the CPE inhibitions were at concentrations and IC_50_ that were consistent with achievable plasma levels at current recommended doses(Erythromycin and pyridoxine) or formulations (Retapamulin) of these drugs. Furthermore, erythromycin, retapamulin and pyridoxine significantly inhibited SARS-CoV-2 M^PRO^ and PP at achievable concentrations and IC_50_. To the best of our knowledge, this is the first study identifying erythromycin and retapamulin as potential therapeutics for COVID-19. Pyridoxine has previously been reported as a useful supplementary therapy for COVID-19 but this was in context unrelated to its antiviral activity^20,21^.

Folic acid significantly inhibited SARS-CoV-2 induced CPE with maximum inhibition of 42% at clinically achievable concentration of 5 μM, thus satisfying our criteria for hits but falling short of the criteria of Severson *et al*^11^. Furthermore, the dose response pattern suggest that lower doses are more effective than higher doses suggesting that higher doses may be toxic to Vero cells in the presence of SARS-CoV-2 viruses because, as reported above, the tested doses of the drug (alone) were not found to be toxic to Vero-cells in the cell toxicity study. This pattern of lower dose being more effective is also observed in Folic acid’s significant inhibition of SARS-CoV-2 M^PRO^. The immediate implication of these findings is uncertain beyond the desirable recommendation that lower doses of folic acid should be the target of future possible utilization of Folic acid in SARS-CoV-2 infections. This is contrary to the suggestion of Asad *et al*^22^ that higher dose of folic acid would be beneficial in COVID-19; though the suggestion was based on studies that examined the inhibition of Spike protein by folic acid^23^. Folic acid also significantly inhibits SARS-CoV-2 Papain like protease and at concentrations and IC_50_ that are also achievable at routine therapeutic doses. Indeed, previous molecular docking studies by our group^4^ and others^24^ suggest Folic acid as potential therapeutic in COVID-19. Wet laboratory studies further suggest that Folic acid inhibits SARS-COV-2 nucleocapsid protein^25^ and inhibits cell invasion by SARS-CoV-2 by methylating ACE2^26^. The relationship between a particular enzyme inhibition and inhibition of CPE is therefore not immediately certain. Perhaps, combination effect may be an explanation.

Ivermectin significantly inhibited SARS-CoV-2 M^PRO^ and PP with either greater or equivalent percentage inhibition compared to other drugs tested. Nonetheless, ivermectin failed to significantly inhibit SARS-CoV-2 induced CPE. These findings may partly explain the lack of clinical efficacy of ivermectin in various clinical trials^27–29^ despite the fact that at very high doses(1,200μg/Kg), started early in the disease ^30^, ivermectin potently inhibit viral replication with up to 5000 fold reduction in viral load^13^. In any case, it is important to note that the concentrations of ivermectin under consideration are not achievable clinically at current recommended doses of <200μg/Kg^14,15,31^. Nonetheless, our findings further point towards CPE and/or cell based assays as, perhaps, more clinically relevant endpoints in screening therapeutics against SARS-CoV-2^5,11,32^.

### Limitations of the study

A limitation of this study is that a limited dose span and restricted dose escalations were used which did not allow for the exploration of the full range of effects of the drugs. However, the restrictions were important in the presence of usual budget constraints, and to focus on achievable concentrations in human plasma because off-label use rather than expanded label was the theoretical framework of this study. Such full-range dose studies will still need to be done if expanded drug labelling that may involve alternative delivery systems are under consideration. Another limitation is that the studies were conducted between three different collaborating laboratories that may have different laboratory fidelities. Nonetheless, this was a pre-identified challenge and steps were taken to harmonize workflow between laboratories and thus enhance data integrity. Even then, no experiment conducted in one laboratory was re-run in another collaborating laboratory. Also, although we estimated IC_50_ in this study, the point estimates should be interpreted with caution because of the possibility of non-monotonicity in the response, a known drawback of IC_50_/EC_50_ estimations^33^(for example, the CPE and M^PRO^ inhibitions of retapamulin appears ‘n’ shaped and ‘u’ shaped respectively)(Figures 1 and 3). The 95% CI estimates of the IC_50_ provide some reassurance, but again these were estimated by using the accepted method of inputting unknown data from standard curves^34,35^. Nonetheless, the strengths of the study are the use of validated cell-based assays, ease of running the experiments in moderately equipped laboratories (International Standard Organization Level 2^36^) and the detection of acceptable level of effects at therapeutic concentrations of the drugs.

### Conclusion

In this study, we identified erythromycin, retapamulin, pyridoxine and folic acid as potential therapeutic agents for COVID-19 and provided evidence that ivermectin may not be effective. Because full or close to full effect (100% inhibition) is an ideal target of drug therapy and none of the drugs achieved this at therapeutic doses, combination therapy is recommended though synergism may not be guaranteed. Such combinations may be evaluated in a randomized control trial using the basket or umbrella design^37^. Such a trial will need careful design and funding consideration. Meanwhile, we consider that the evidence provided by this study is sufficient for consideration of off-label use of these drugs in COVID-19 situations; given that the evidence is consistent and comparable with those available for drugs currently on clinical trials^38,39^ and probably superior to those currently on off-label use^40,41^. We also recommend that all off-label prescriptions, while maintaining standard ethical requirements, should include robust and objective documentation of patients’ status and drug dosing to provide *preliminary insight* into the effectiveness of the drugs. Such *Preliminary Insight documentation* should be routine in off-label prescriptions but may not and cannot replace randomized trials. In this regard, risk of abuse of documented off-label prescription, like using it as convenient alternative to clinical trials, should be recognized and mitigated^41^.

